# Lack of sex differences in gonadotropin-releasing hormone (GnRH) neuron potassium currents and excitability

**DOI:** 10.1101/2021.03.24.436851

**Authors:** R. Anthony DeFazio, Suzanne M. Moenter

**Affiliations:** Departments of Molecular & Integrative Physiology, University of Michigan, Ann Arbor, MI; Obstetrics and Gynecology, University of Michigan, Ann Arbor, MI; Internal Medicine, University of Michigan, Ann Arbor, MI

## Abstract

Gonadotropin-releasing hormone (GnRH) drives pituitary secretion of luteinizing hormone (LH) and follicle-stimulating hormone, which in turn regulate gonadal functions including steroidogenesis. The pattern of GnRH release and thus fertility depend on gonadal steroid feedback. Under homeostatic (negative) feedback conditions, removal of the gonads from either females or males increases the amplitude and frequency of GnRH release and alters the long-term firing pattern of these neurons in brain slices. The neurobiological mechanisms intrinsic to GnRH neurons that are altered by homeostatic feedback are not well studied and have not been compared between sexes. During estradiol positive feedback, which is unique to females, there are correlated changes in voltage-gated potassium currents and neuronal excitability. We thus hypothesized these same mechanisms would be engaged in homeostatic negative feedback. Voltage-gated potassium channels play a direct role in setting excitability and action potential properties. Whole-cell patch-clamp recordings of GFP-identified GnRH neurons in brain slices from sham-operated and castrated adult female and male mice were made to assess fast (I_A_) and slow (I_K_) inactivating potassium currents as well as action potential properties. Surprisingly, no changes were observed in most potassium current properties, input resistance or capacitance and this was reflected in a lack of differences in excitability and specific action potential properties. These results support the concept that, in contrast to positive feedback, steroid negative feedback regulation of GnRH neurons in both sexes is likely conveyed to GnRH neurons via mechanisms that do not induce major changes in the biophysical properties of these cells.

**Significance Statement:** The pattern of activity of gonadotropin-releasing hormone (GnRH) neurons is crucial to reproductive success in both males and females. Direct comparison of GnRH neurons from mice of both sexes during negative feedback and after gonadectomy revealed few differences in potassium currents, excitability, and action potential properties. These results support the hypothesis that neurons presynaptic to GnRH neurons communicate negative feedback to these cells in a manner that does not alter their intrinsic biophysical properties.

## Introduction

The episodic release of GnRH from the brain is key to successful reproduction in both sexes. GnRH regulates pituitary release of luteinizing hormone (LH) and follicle-stimulating hormone, which activate gonadal functions including production of sex steroids. These steroids feed back to regulate the pattern of GnRH release and pituitary response to GnRH. Aspects of this feedback system are well established as sexually differentiated, for example estradiol positive feedback is exclusive to females under normal physiologic conditions and induces the preovulatory surge of GnRH (Docke and Dorner, 1965; Sarkar et al., 1976; Moenter et al., 1991). Other aspects appear similar between the sexes, including homeostatic negative feedback that regulates episodic GnRH release and is critical for maintaining most reproductive processes in both males and females. Gonadectomy increases GnRH and LH release (Leipheimer et al., 1985; Levine et al., 1985; Karsch et al., 1987; Caraty and Locatelli, 1988; Condon et al., 1988; Jackson and Kuehl, 2000; Czieselsky et al., 2016). The main circulating sex steroids providing homeostatic feedback are sexually differentiated, specifically estradiol and progesterone in females, depending on stage of the reproductive cycle, and testosterone in males. The differences in circulating steroids are reduced in part in the brain by conversion of testosterone to estradiol (Fisher et al., 1998; Sharma et al., 1999), and the efficacy of androgens and estrogens in eliciting negative feedback in males varies with species (Plant, 1982; Levine and Duffy, 1988; Tilbrook et al., 1999).

Regulation of episodic GnRH release is a key potential intervention point for the very different goals of reversibly inhibiting reproduction for contraception and ameliorating central infertility. Despite a similar hormonal response to the removal of peripheral sex steroid feedback by gonadectomy, whether or not the underlying intrinsic changes in GnRH neurons that lead to these increases are sexually differentiated is unknown. Studies in females have focused on the mechanisms underlying estradiol positive feedback that is critical for inducing ovulation; many of these have compared gonadectomized animals with open feedback loops to those with specific steroid replacement (Wagner et al., 2001; Christian and Moenter, 2010; Dror et al., 2013; Liu et al., 2017; Adams et al., 2018b; Adams et al., 2019; Wang et al., 2019). These studies have revealed changes in both intrinsic GnRH neuron properties and fast synaptic input to these cells. Studies of homeostatic negative feedback suggest that the firing pattern of GnRH neurons varies during the reproductive cycle in females and when steroid feedback is disrupted by gonadectomy in both sexes (Pielecka and Moenter, 2006; Pielecka et al., 2006; Silveira et al., 2017). There are no direct comparisons, however, of GnRH neuron intrinsic properties between intact males and females or how these are affected by gonadectomy.

A basic measure of the intrinsic biophysical properties of neurons is their excitability, defined as the number of action potentials generated in response to varying current injections. In females, this varies with cycle stage and with induction of daily LH surges by estradiol in ovariectomized mice (Adams et al., 2018b; Adams et al., 2019) but this parameter has not been characterized in males. Voltage-gated potassium channels are widely recognized as regulators of excitability (Johnston et al., 2010) and have been targeted clinically. For example, blocking a fraction of potassium channels with 4-aminopyridine (4AP) increases the excitability of motor neurons in amyotropic lateral sclerosis and multiple sclerosis and provides relief from the symptoms of these diseases (Bakirtzis et al., 2018; Peikert et al., 2019). In many neurons, 4AP increases input resistance and diminishes the ability of voltage-gated potassium channels to blunt action potential firing and/or the membrane potential changes in response to synaptic inputs (Hoffman, 2013; DeFazio et al., 2019). Here we tested the hypotheses that voltage-gated potassium currents are reduced and excitability of GnRH neurons increased by gonadectomy in both sexes.

## Materials and Methods

All chemicals were acquired from Sigma-Aldrich (St. Louis, MO, USA) unless noted.

### Animals

Mice expressing GFP under control of the GnRH promoter (Tg(GnRH1-EGFP)51Sumo MGI:6158457, GnRH-GFP mice, JAX 033639) were propagated in our colony. All mice had *ad libitum* access to Harlan 2916 chow and water and were held at 21-23 °C on a 14L:10D light cycle with lights on at 0300 eastern standard time. Adult mice 82-146 days old were ovariectomized (OVX, females), orchidectomized (ORX, males) or sham operated under isoflurane anesthesia with bupivacaine as a local analgesic. Studies were performed 5-7 days after surgery. In sham-operated females, estrous cycles were monitored by vaginal cytology for at least 10 days before experiments and diestrous mice were selected. Body, uterine and seminal vesicle mass were recorded at the time of brain slice preparation to verify steroid status. The Institutional Animal Care and Use Committee of the University of Michigan approved all procedures.

### Brain slice preparation

All extracellular solutions were bubbled with 95% O_2_/5% CO_2_ throughout the experiments and for at least 30 min before exposure to tissue. Coronal brain slices (300 µm) containing the preoptic area (POA) were prepared with a Leica VT1200S (Leica Biosystems) using modifications of previously described methods (DeFazio and Moenter, 2002). Brain slices were obtained between 0730-0830 EST; recordings obtained between 0830-1430 EST. The brain was rapidly removed and placed in ice-cold sucrose saline solution containing (in mM): 250 sucrose, 3.5 KCl, 26 NaHCO_3_, 10 D-glucose, 1.25 Na_2_HPO_4_, 1.2 MgSO_4_, and 3.8 MgCl_2_ (350 mOsm). Slices were incubated for 30 min at room temperature (∼21–23°C) in 50% sucrose saline and 50% artificial cerebrospinal fluid (ACSF, containing (in mM): 135 NaCl, 3.5 KCl, 26 NaHCO_3_, 10 D-glucose, 1.25 Na_2_HPO_4_, 1.2 MgSO_4_, 2.5 CaCl_2,_ 315 mOsm, pH 7.4). Slices were then transferred to 100% ACSF solution at room temperature for 0.5–5h before recording. One to three recordings were obtained per mouse with a minimum of 5 mice studied per group; variation within a mouse was not less than that within a group.

### Recording solutions and data acquisition

The pipette solution consisted of (in mM): 125 K gluconate, 20 KCl, 10 HEPES, 5 EGTA, 0.1 CaCl_2_, 4 MgATP and 0.4 NaGTP, 305 mOsm, Ph 7.2 with NaOH; this solution is based on the native intracellular chloride concentration in GnRH neurons determined using gramicidin perforated-patch recordings (DeFazio et al., 2002). A 14.5 mV liquid junction potential was negated before each recording (Barry, 1994). During all recordings, slices were continuously superfused at 2 ml/min with carboxygenated ACSF kept at 30-31°C with an inline-heating unit (Warner Instruments). GFP-positive cells were visualized with a combination of infrared differential interference contrast and fluorescence microscopy on an Olympus BX50WI microscope. Recordings were made with an EPC-10 patch clamp amplifier and a computer running PATCHMASTER software (HEKA Elektronik). For current-clamp experiments, membrane voltage was acquired at 20 kHz and filtered at 10 kHz; for voltage-clamp experiments, current was acquired at 10 kHz and filtered at 5 kHz. Input resistance, series resistance, baseline current, and capacitance were monitored throughout experiments from the membrane current response to a 20 ms, 5 mV hyperpolarizing voltage step from −65 mV to monitor recording quality. All recordings with input resistances (R_in_) <0.5 GΩ, series resistances (R_s_) >20 MΩ, or unstable membrane capacitance (C_m_) were rejected. Results did not depend on anatomical location of GnRH neurons within the POA.

### Experimental Design

Potassium currents, action potential properties and passive properties were characterized in GFP-identified GnRH neurons in the preoptic area of brain slices prepared from adult gonad intact and castrated mice of both sexes. Castrated mice were studied 5-7 days post surgery; intact females were in diestrus.

### Voltage-gated potassium current characterization

Potassium currents were isolated pharmacologically during whole-cell voltage-clamp recordings by blocking fast sodium and calcium channels as well as ionotropic receptors for GABA and glutamate (2 µM tetrodotoxin (TTX, Tocris), 300 µM NiCl_2_, 20 µM D-APV (Tocris), 10 µM CNQX, and 100 µM picrotoxin). Membrane potential was held at −65 mV between voltage-clamp protocols. Series resistance was monitored before compensation using the current response to a 5 mV hyperpolarizing step. Cells with R_s_>20 MΩ without compensation were discarded. Two potassium currents, I_A_ and I_K_, were distinguished based on voltage dependence and time course. I_A_ is a typical, rapidly-inactivating potassium current that can be activated at membrane potentials that are hyperpolarized to the threshold for action potential firing. I_K_ also displays voltage-dependent inactivation but this is restricted to more depolarized potentials and has a slower time course of inactivation.

### Activation and inactivation of IA

Preliminary studies on I_A_ showed that a 500 ms prepulse at −40 mV induced full inactivation, and a prepulse at −100 mV for 500 ms completely removed inactivation. These two prepulses were combined with a series of voltage steps to isolate and characterize I_A_. To measure total potassium current, a 500 ms prepulse at −100 mV was first applied to remove the inactivation of the fast transient component, followed by a family of voltage steps (500 ms, 10 mV intervals) from −100 mV to +50 mV, and then a final step to −10 mV for 100 ms. To inactivate I_A_, the same step protocol was applied but with the 500 ms prepulse set at −40 mV to inactivate the fast component while leaving I_K_ mostly unchanged attributable to its very slow inactivation (see below). The fast I_A_ component was then isolated by subtracting the family of currents obtained with the −40 mV prepulse from that obtained with the −100 mV prepulse. Activation of I_A_was quantified by measuring the peak current reached during each voltage step in the family (see Analysis section below). Inactivation of I_A_ was estimated from the peak current during the final step to −10 mV that followed the family of voltage steps. All protocols for I_A_ analysis were leak subtracted using the online -P/8 (average of 8 sweeps, 1/8 the size of the voltage step in the opposite direction, from a baseline potential of −65 mV) (Bezanilla and Armstrong, 1977).

Time course of inactivation of I_A_ was characterized by stepping the membrane potential to −100 mV for 500 ms to remove inactivation, then stepping to the inactivation potential (−40 mV) for 0, 0.5, 1, 2, 4, 8, 16, 32, 64, 128, 256, 512, 1024 ms, followed by a test pulse at −10 mV to assess the peak current. The non-inactivating component during the test pulse after 1024 ms inactivation was subtracted from each of the other traces to isolate the transient current. Similarly, the time course of recovery from inactivation of I_A_ was characterized by stepping the membrane potential to −40 mV for 500 ms to fully inactivate I_A_, and then stepping to −100 mV for the durations above, followed by a test pulse at −10 mV to assess the peak current. The non-inactivating component at 0 ms recovery was subtracted from each of the other traces to isolate the transient current.

To study I_K_, a separate set of recordings was required because of the slow inactivation of this component. Indeed full inactivation and recovery required >10 s at +50 mV and −100 mV, respectively. Cells did not remain stable upon repeated exposure to these potentials, thus more moderate potentials were used to permit estimation of these properties within the command potentials and durations the cells tolerated.

### Activation and inactivation of I_K_

To quantify activation, a double prepulse protocol was used: an initial prepulse to −75 mV for 10 s was used to remove inactivation from I_K_, then a second prepulse to −50 mV for 1 s was used to inactivate I_A_. After these two prepulses, test pulses of 10 s from −50 to +50 mV in 10 mV increments were used to measure the peak I_K_ current. A final step to +50 mV for 100 ms was used following each test pulse to measure inactivation of I_K_. The long duration voltage steps required to characterize this current makes traditional online leak subtraction unrealistic. For the activation of I_K_, leak currents were subtracted offline using the approach of (Kimm and Bean, 2014). The shape of the leak response was acquired from the average of 16 steps of −5 mV for 50 ms from the holding potential of −65 mV. Offline, the average was scaled by the voltage difference and subtracted from the portion of the raw current sweep containing the activation test pulses. The leak steps were run immediately before each voltage-clamp protocol. Prior to each leak acquisition, passive properties were recorded and both slow capacitance and series resistance compensation (50-70%) adjusted.

Time course of inactivation of I_K_ was characterized by first removing inactivation by holding at - 75 mV for 10s, then I_A_ was inactivated by stepping to −50mV for 1 s. This was followed by a variable duration inactivation pulse (0.1, 0.21, 0.42, 0.83, 1.64, 3.25, 6.46, 12.87, 25.68, 51.29 s) at −30 mV, followed by a test pulse at +50 mV. Because inactivation was incomplete even after 51.29 s at −30 mV, this remaining current was not subtracted and the inactivation graphs level off at ∼30%. Recovery from inactivation for I_K_ was studied using an inactivating prepulse of +50 mV for 10 s, followed by a variable duration recovery pulse at −80 mV (0.01, 0.03, 0.06, 0.11, 0.2, 0.37, 0.7, 1.35, 2.64, 5.21, 10.34, 20.59 s), a brief pulse to inactivate I_A_ (−50 mV for 1 s), and a test pulse at +50 mV for 100 ms. The peak current was plotted as a function of the recovery prepulse duration. Since inactivation was incomplete, the non-inactivated current was not subtracted (recovery graphs start at ∼30%).

### Analysis

Current density was calculated by dividing current by capacitance for each cell. To assess the voltage-dependence of activation, both I_A_ and I_K_ were divided by the driving force derived from the Goldman-Hodgkin-Katz (GHK) current equation (Clay, 2000, 2009). To estimate V_0.5activation_, I_A_ and I_K_ were divided by the GHK driving force, normalized to the maximum value, plotted as a function of step potential and fit with the Boltzmann equation:

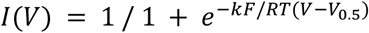

V is the command potential of the step, V_0.5_ is the potential at half maximum, and k is the “slope factor” (k has no units attributable to the F/RT factor). Voltage dependence of inactivation was characterized using the same equation to fit the normalized current measured during the inactivation test pulse.

### Action potential properties

To characterize action potential properties, current-clamp recordings were obtained in the presence of antagonists of receptors for fast synaptic transmission (20 µM D-APV, 10 µM CNQX, 100 µM picrotoxin). Cells were maintained at −65 mV by current injection (<50 pA) in current-clamp after bridge compensation of series resistance by at least 95%. Current steps (5 pA increments, 500 ms, −10 pA to +50 pA, 2.5 s interval near −65 mV between sweeps) were delivered to test the membrane potential response. The first current step to display an action potential was defined as the rheobase and the first spike analyzed in detail. Action potential threshold was defined as the potential at which the membrane potential slope exceeded 1 V/s. Action potential latency was defined as the time from start of the current injection to threshold. Rate of rise was defined as the maximum of the voltage trace derivative from threshold to peak. Full width of the action potential at half maximum (FWHM), and afterhyperpolarization (AHP) time and amplitude were measured relative to threshold.

### Statistics

Data are reported as mean ± SEM, with individual values shown where practical. Statistical comparisons were made using Prism 9 (GraphPad Software). Data were tested for normal distribution with Shapiro-Wilk. Tests were chosen appropriate for the experimental design and data distribution as specified in the results. For two-way ANOVA, the Bonferroni *post hoc* is considered sufficiently robust to use with non-normally distributed data (Underwood, 1996). Significance was set at p<0.05.

## Results

### Verification of gonadectomy

Short-term gonadectomy had no effect on body mass within either sex (females n=12 sham diestrus 21.4±0.6 g, n=14 OVX 21.8±0.3 g; males n=13 sham 26.1±0.7 g, n=13 ORX 25.4±0.6 g, two-way ANOVA gonadal status: F(1, 48) = 0.07108; P=0.7909). As expected, males were heavier than females (two-way ANOVA sex: F(1, 48) = 53.80; P<0.0001). In females, OVX reduced uterine mass (sham diestrus 63.4±5.0 mg, OVX 29.6±1.3 mg, unpaired two-tailed Student’s t test: t=7.001, df=24, P<0.0001). In males, ORX reduced seminal vesicle mass (sham 217.2±11.6 mg, ORX 56.6±4.4 mg, unpaired two-tailed Student’s t test: t=12.97, df=24, P<0.0001).

### Passive properties of GnRH neurons do not vary with sex or gonadal status under conditions used to characterize potassium currents

Critical to evaluating of current properties among treatments is comparison of similar quality recordings. No difference was observed among groups for uncompensated series resistance (Figure 1A). Nor were any differences in the passive cellular properties of input resistance (Figure 1B), capacitance (Figure 1C) or holding current (Figure 1D) attributable to either sex or gonadal status. These latter measures offer a combined view of recording quality as well as insight into potential biological changes, such as cell size or membrane conductance (two-way ANOVA, Table 1).

**Table 1.** Statistical parameters from two-way ANOVA for passive properties from potassium current recordings (Figure 1)

| Property | Sex | Gonadal Status | Interaction |
| --- | --- | --- | --- |
| Series resistance | F(1, 56) = 1.512;<br>P=0.2240 | F(1, 56) = 3.308;<br>P=0.0743 | F(1, 56) = 0.1171;<br>P=0.7335 |
| Input resistance | F(1, 56) = 0.03673;<br>P=0.8487 | F(1, 56) = 0.3186;<br>P=0.5747 | F(1, 56) = 0.4542;<br>P=0.5031 |
| Capacitance | F(1, 56) = 0.1963;<br>P=0.6594 | F(1, 56) = 2.540;<br>P=0.1166 | F(1, 56) = 0.08991;<br>P=0.7654 |
| Holding current | F(1, 56) = 2.592;<br>P=0.1130 | F(1, 56) = 0.6304;<br>P=0.4306 | F(1, 56) = 1.496;<br>P=0.2263 |

**Figure 1.**
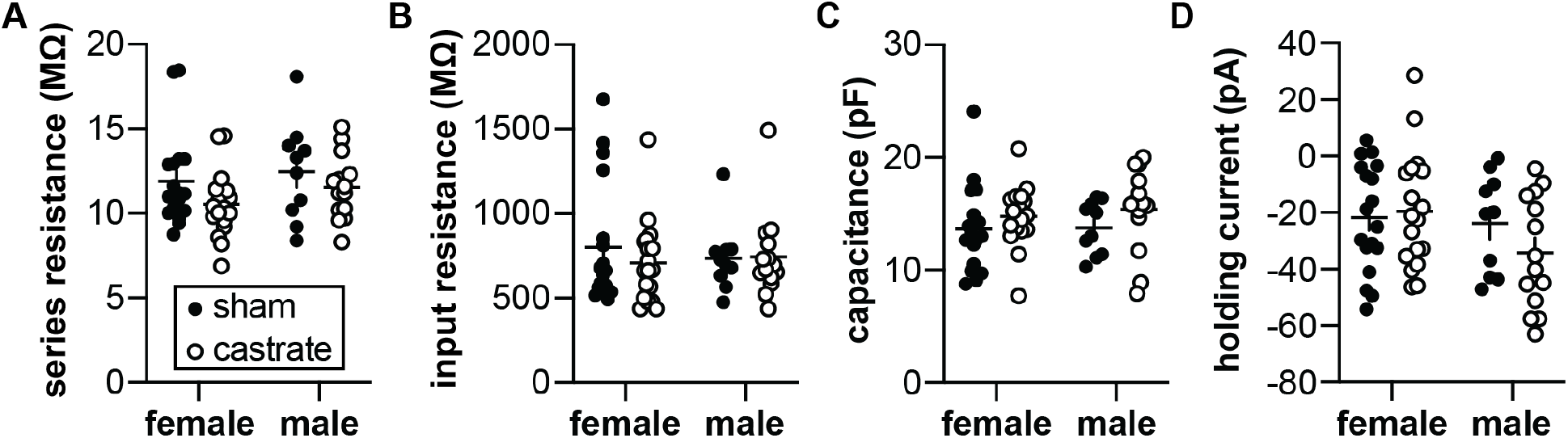
Passive properties of GnRH neurons in the potassium current experiments. No differences attributable to sex or gonadal status were detected in (A) series resistance, (B) input resistance, (C) capacitance or (D) holding current using two-way ANOVA (Table 1).

### Neither sex nor gonadal status affect voltage-gated potassium currents of GnRH neurons

The output of GnRH neurons in terms of spontaneous firing pattern and hormone release is increased by removal of homeostatic gonadal steroid feedback accomplished via gonadectomy. We hypothesized that removing gonadal steroid feedback would reduce voltage-gated potassium currents. Two components of this current were examined, a fast I_A_-like current and a slowly-inactivating I_K_-like current. No measured property of either current differed with sex or gonadal status (Figures 2, 3; Tables 3, 4). Only the parameters calculated from the Boltzmann fit, specifically the inactivation V0.5 and slope factor for I_A_ and both inactivation and activation slope factors for I_K_ exhibited weakly significant P values.

**Figure 2.**
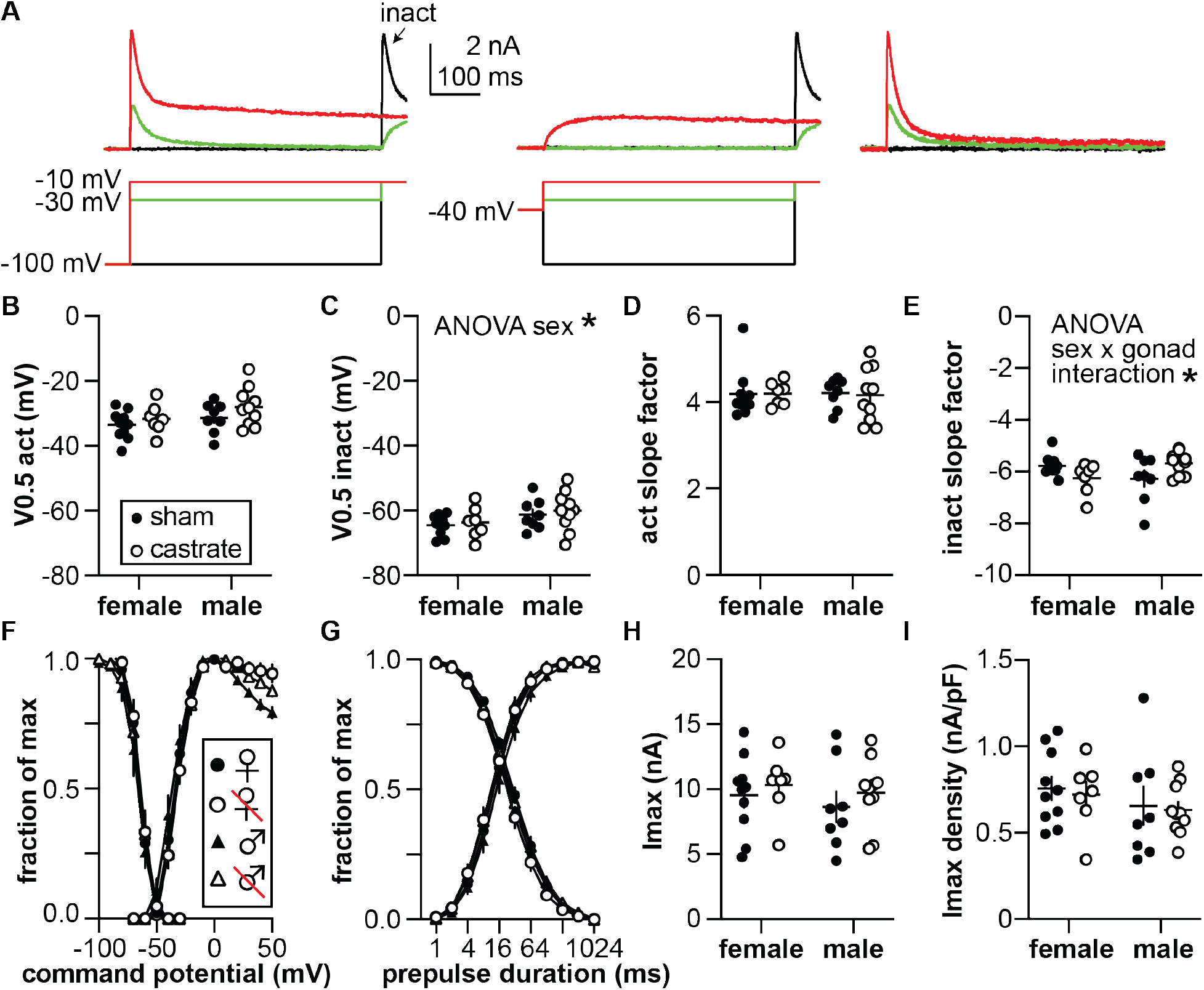
Characterization of the I_A_ potassium current. (A) Representative traces illustrating the isolation of the rapidly inactivating I_A_ (top) and the voltage protocols used (bottom). The 500 ms 100 mV prepulse was truncated for illustration. The arrow labeled “inact” indicates the −10 mV test pulse at the end of the activation voltage family used to calculate the voltage dependence of inactivation. Only three steps from the voltage family from −100 to +50 mV are shown for clarity. The right panel shows I_A_ isolated by subtracting the −40 mV prepulse traces in the middle panel from the −100 mV prepulse traces in the left panel. (B, C) membrane potential at which half of the current is activated (V0.5act) (B) or inactivated (V0.5inact) (C). (D, E) activation (act) (D) and inactivation (inact) (E) slope factors. (F) Voltage dependence of activation and inactivation, normalized by maximum conductance and maximum current respectively. (G) Time course of recovery and inactivation, normalized by maximum current. (H, I) Maximum current (Imax) (H) and current density (I). Statistical parameters are in Table 2.

**Table 2.** Two-way ANOVA statistical parameters for I_A_. Bold indicates P<0.05.

| Property | Sex | Gonadal Status | Interaction |
| --- | --- | --- | --- |
| V0.5 activation | F(1, 31) = 2.921;<br>P=0.0974 | F(1, 31) = 2.361;<br>P=0.1345 | F(1, 31) = 0.1956;<br>P=0.6614 |
| V0.5 inactivation | **F(1, 31) = 4.473;**<br>**P=0.0426** | F(1, 31) = 0.4313;<br>P=0.5162 | F(1, 31) = 0.008208;<br>P=0.9284 |
| activation slope factor | F(1, 31) = 0.004146;<br>P=0.9491 | F(1, 31) = 0.01815;<br>P=0.8937 | F(1, 31) = 0.02354;<br>P=0.8791 |
| inactivation slope factor | F(1, 31) = 0.02076;<br>P=0.8864 | F(1, 31) = 0.07103;<br>P=0.7916 | **F(1, 31) = 6.872;**<br>**P=0.0134** |
| I<sub>max</sub> | F(1, 31) = 0.5863;<br>P=0.4496 | F(1, 31) = 0.9055;<br>P=0.3487 | F(1, 31) = 0.02845;<br>P=0.8671 |
| I<sub>max</sub> density | F(1, 31) = 1.576;<br>P=0.2188 | F(1, 31) = 0.1613;<br>P=0.6907 | F(1, 31) = 0.006205;<br>P=0.9377 |

**Figure 3.**
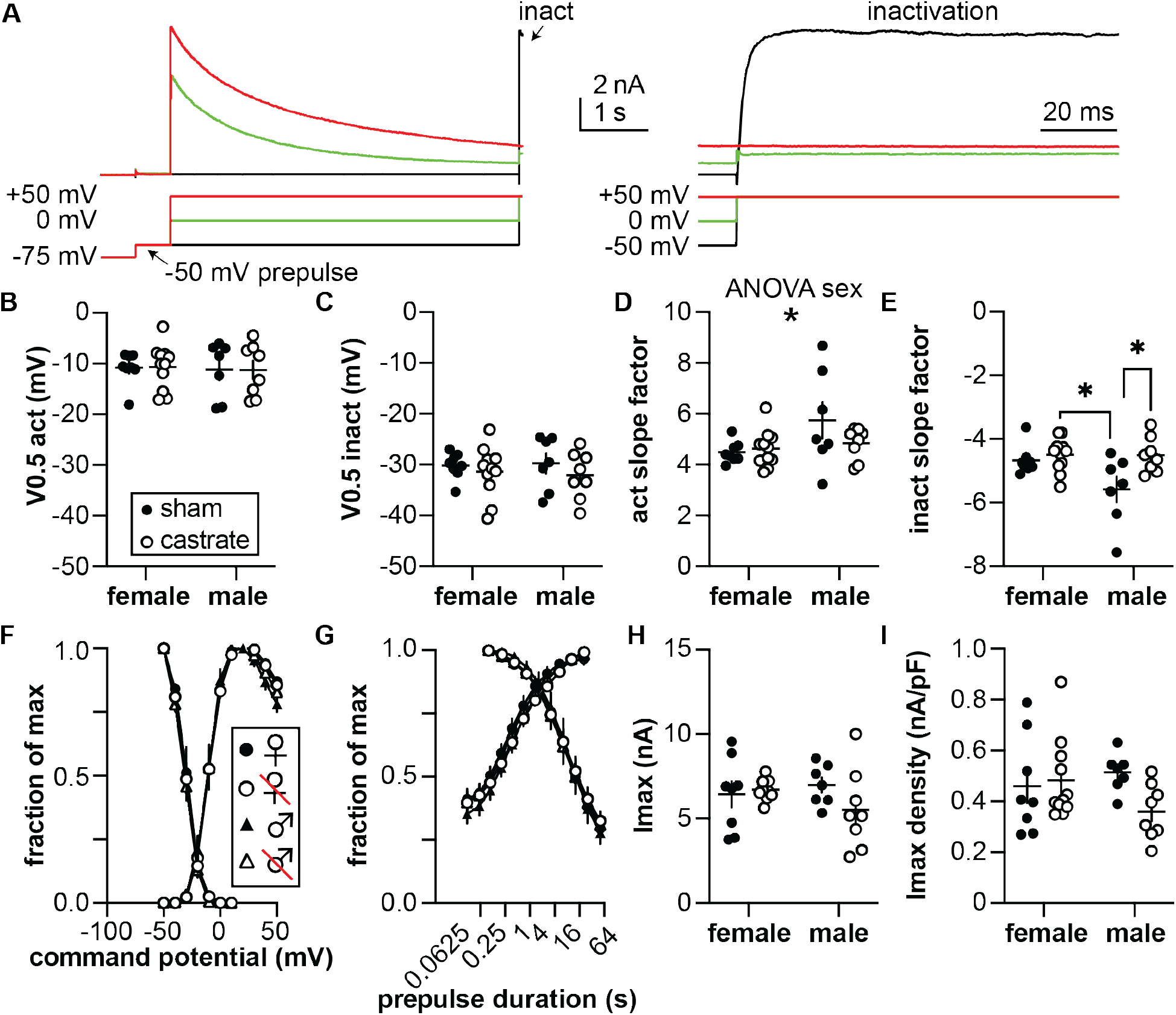
Characterization of the I_k_ potassium current. (A) Representative traces illustrating the activation and inactivation of I_K_, (top) and the voltage protocols used (bottom). The 10 s −75 mV prepulse was truncated in the left panel for illustration. The arrow labeled “inact” on the left panel indicates the region expanded on the right. Only three steps of the voltage family from − 100 to +50 mV are shown for clarity. (B, C) membrane potential at which half of the current is activated (V0.5act) (B) or inactivated (V0.5inact) (C). (D, E) activation (act) (D) and inactivation (inact) (E) slope factors. (F) Voltage dependence of activation and inactivation, normalized by maximum conductance and maximum current respectively. (G) Time course of recovery and inactivation, normalized by maximum current. (H, I) Maximum current (Imax) (H) and current density (I). Statistical parameters are in Table 3.

**Table 3.** Two-way ANOVA statistical parameters for I_K_. Bold indicates P<0.05.

| Property | Sex | Gonadal Status | Interaction |
| --- | --- | --- | --- |
| V0.5 activation | F(1, 30) = 0.09013;<br>P=0.7661 | F(1, 30) = 0.0003841;<br>P=0.9845 | F(1, 30) = 0.006304;<br>P=0.9372 |
| V0.5 inactivation | F(1, 30) = 0.006266;<br>P=0.9374 | F(1, 30) = 1.247;<br>P=0.2731 | F(1, 30) = 0.1239;<br>P=0.7273 |
| activation slope factor | **F(1, 30) = 4.294;**<br>**P=0.0469** | F(1, 30) = 1.170;<br>P=0.2880 | F(1, 30) = 2.197;<br>P=0.1487 |
| inactivation slope factor | F(1, 30) = 3.971;<br>P=0.0554 | **F(1, 30) = 7.378;**<br>**P=0.0109** | F(1, 30) = 3.864;<br>P=0.0587 |
| I<sub>max</sub> | F(1, 30) = 0.2979;<br>P=0.5892 | F(1, 30) = 1.019;<br>P=0.3208 | F(1, 30) = 2.259;<br>P=0.1433 |
| I<sub>max</sub> density | F(1, 30) = 0.4582;<br>P=0.5036 | F(1, 30) = 1.725;<br>P=0.1990 | F(1, 30) = 2.988;<br>P=0.0942 |

**Table 4.** Two-way ANOVA statistical parameters for passive properties from action potential recordings (Figure 5). Bold indicates P<0.05.

| Property | Sex | Gonadal status | Interaction |
| --- | --- | --- | --- |
| Series resistance | $F(1, 37) = 0.004601$;<br>$P = 0.9463$ | $F(1, 37) = 0.3899$;<br>$P = 0.5362$ | **$F(1, 37) = 4.366$;**<br>**$P = 0.0436$** |
| Input resistance | $F(1, 37) = 0.03866$;<br>$P = 0.8452$ | $F(1, 37) = 0.1832$;<br>$P = 0.6712$ | $F(1, 37) = 0.4448$;<br>$P = 0.5089$ |
| Capacitance | $F(1, 37) = 0.1299$;<br>$P = 0.7205$ | $F(1, 37) = 1.521$;<br>$P = 0.2253$ | $F(1, 37) = 1.343$;<br>$P = 0.2539$ |
| Holding current | $F(1, 37) = 1.555$;<br>$P = 0.2202$ | $F(1, 37) = 0.5610$;<br>$P = 0.4586$ | $F(1, 37) = 0.2843$;<br>$P = 0.5971$ |

### Passive properties of GnRH neurons do not vary with sex or gonadal status under conditions used to characterize action potentials

To record action potential properties, the blockers of voltage-gated calcium and sodium channels needed to isolate potassium currents were omitted; this may alter the passive properties of the cells, thus these parameters are reported separately for the two types of recordings. There was a weak interaction between sex and gonadal status for uncompensated series resistance (Figure 4A, P=0.0436). This difference is unlikely to account for a difference in measured values as bridge balance was in effect to compensate series resistance by >95% during action potential recordings. As was observed for recordings of potassium currents, there were no differences in the passive cellular properties of input resistance (Figure 4B), capacitance (Figure 4C) or holding current (Figure 4D) attributable to either sex or gonadal status.

**Figure 4.**
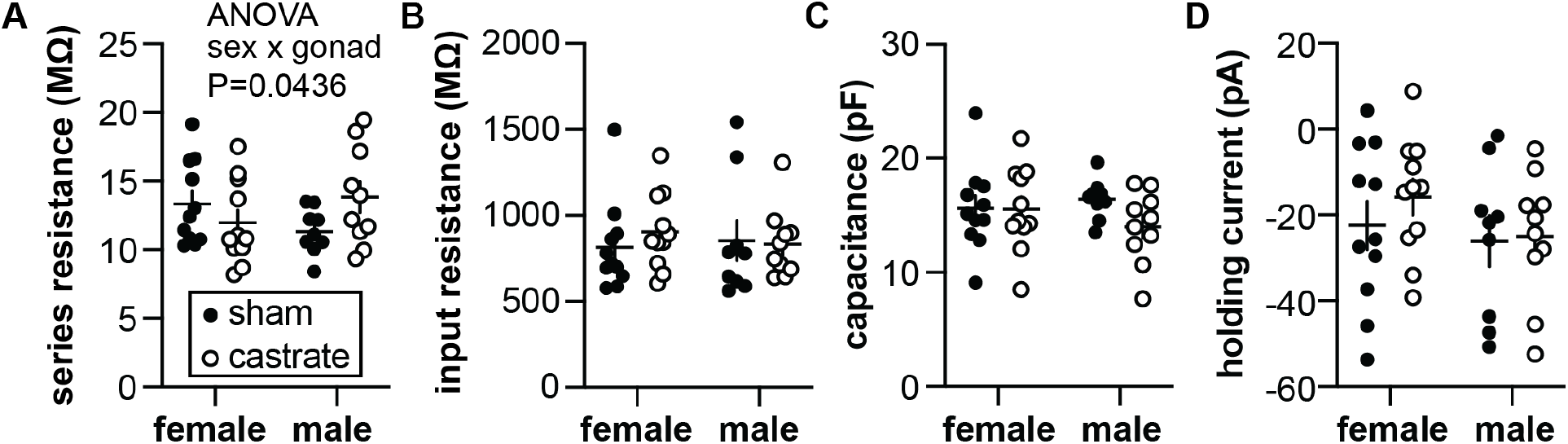
Passive properties of recordings for the action potential study. (A) series resistance (no post hoc comparisons were significant); (B) input resistance; (C) capacitance; (D) holding current at −65 mV.

### Neither excitability nor first action potential properties are altered by sex or gonadal status

Current-clamp recordings of action potential properties were used to assess possible general differences in the intrinsic properties of GnRH neurons in the experimental groups (Figure 5). No differences were observed in the firing response to current injection attributable to sex or gonadal status (three-way repeated-measures ANOVA, Table 5). Post-analysis consolidation of these data to examine effects of either sex or gonadal status by two-way repeated measures ANOVA revealed a mild interaction between sex and current injection, but no post hoc significance (Bonferroni); there was no effect of gonadectomy. There were also no differences observed in any action potential property examined (two-way ANOVA, Table 6). Together these observations suggest that homeostatic feedback does not induce major changes in action potentials of GnRH neurons.

**Table 5.** Three-way repeated measures parameters for action potential firing as a function of current injection. Bold indicates P<0.05.

|  | 3-way | 2-way (sex consolidated) | 2-way (gonadectomized (GDX) consolidated) |
| --- | --- | --- | --- |
| current step (pA) | **$F(2.554, 91.96) = 641.8$;**<br>**$P < 0.0001$** | **$F(10, 380) = 653$;**<br>**$P < 0.0001$** | **$F(10, 380) = 636$;**<br>**$P < 0.0001$** |
| sex | $F(1, 36) = 2.447$;<br>$P = 0.1265$ | $F(1, 38) = 2.6$;<br>$P = 0.1119$ | |
| GDX | $F(1, 36) = 0.4993$;<br>$P = 0.4844$ | | $F(1, 38) = 0.64$;<br>$P = 0.4271$ |
| pA x sex | $F(10, 360) = 1.758$;<br>$P = 0.0668$ | **$F(10, 380) = 1.9$;**<br>**$P = 0.0477$** | |
| pA x GDX | $F(10, 360) = 0.7544$;<br>$P = 0.6728$ | | $F(10, 380) = 0.88$;<br>$P = 0.5513$ |
| sex x GDX | $F(1, 36) = 0.2066$;<br>$P = 0.6522$ | | |
| pA x sex x GDX | $F(10, 360) = 0.5547$;<br>$P = 0.8503$ | | |

**Table 6.** Two-way ANOVA parameters for action potential properties

| **Property** | **Sex** | **Gonadal Status** | **Interaction** |
| --- | --- | --- | --- |
| max frequency | F(1, 37) = 1.463;<br>P=0.2341 | F(1, 37) = 0.7278;<br>P=0.3991 | F(1, 37) = 0.6354;<br>P=0.4305 |
| baseline | F(1, 37) = 0.6163;<br>P=0.4374 | F(1, 37) = 0.1470;<br>P=0.7036 | F(1, 37) = 0.2182;<br>P=0.6431 |
| threshold | F(1, 37) = 0.7224;<br>P=0.4008 | F(1, 37) = 0.001493;<br>P=0.9694 | F(1, 37) = 0.002371;<br>P=0.9614 |
| rheobase | F(1, 37) = 2.778;<br>P=0.1040 | F(1, 37) = 7.015e-005;<br>P=0.9934 | F(1, 37) = 0.8335;<br>P=0.3672 |
| AP latency | F(1, 37) = 0.1120;<br>P=0.7398 | F(1, 37) = 0.1392;<br>P=0.7112 | F(1, 37) = 0.1060;<br>P=0.7466 |
| AP amplitude | F(1, 37) = 1.173;<br>P=0.2858 | F(1, 37) = 0.9688;<br>P=0.3314 | F(1, 37) = 0.4683;<br>P=0.4980 |
| AP FWHM | F(1, 37) = 2.817;<br>P=0.1017 | F(1, 37) = 3.069;<br>P=0.0881 | F(1, 37) = 2.858;<br>P=0.0993 |
| AP rate of rise | F(1, 37) = 0.07317;<br>P=0.7883 | F(1, 37) = 0.2856;<br>P=0.5962 | F(1, 37) = 0.2910;<br>P=0.5928 |
| AHP amp | F(1, 37) = 3.374;<br>P=0.0743 | F(1, 37) = 0.3165;<br>P=0.5771 | F(1, 37) = 0.1254;<br>P=0.7253 |

**Figure 5.**
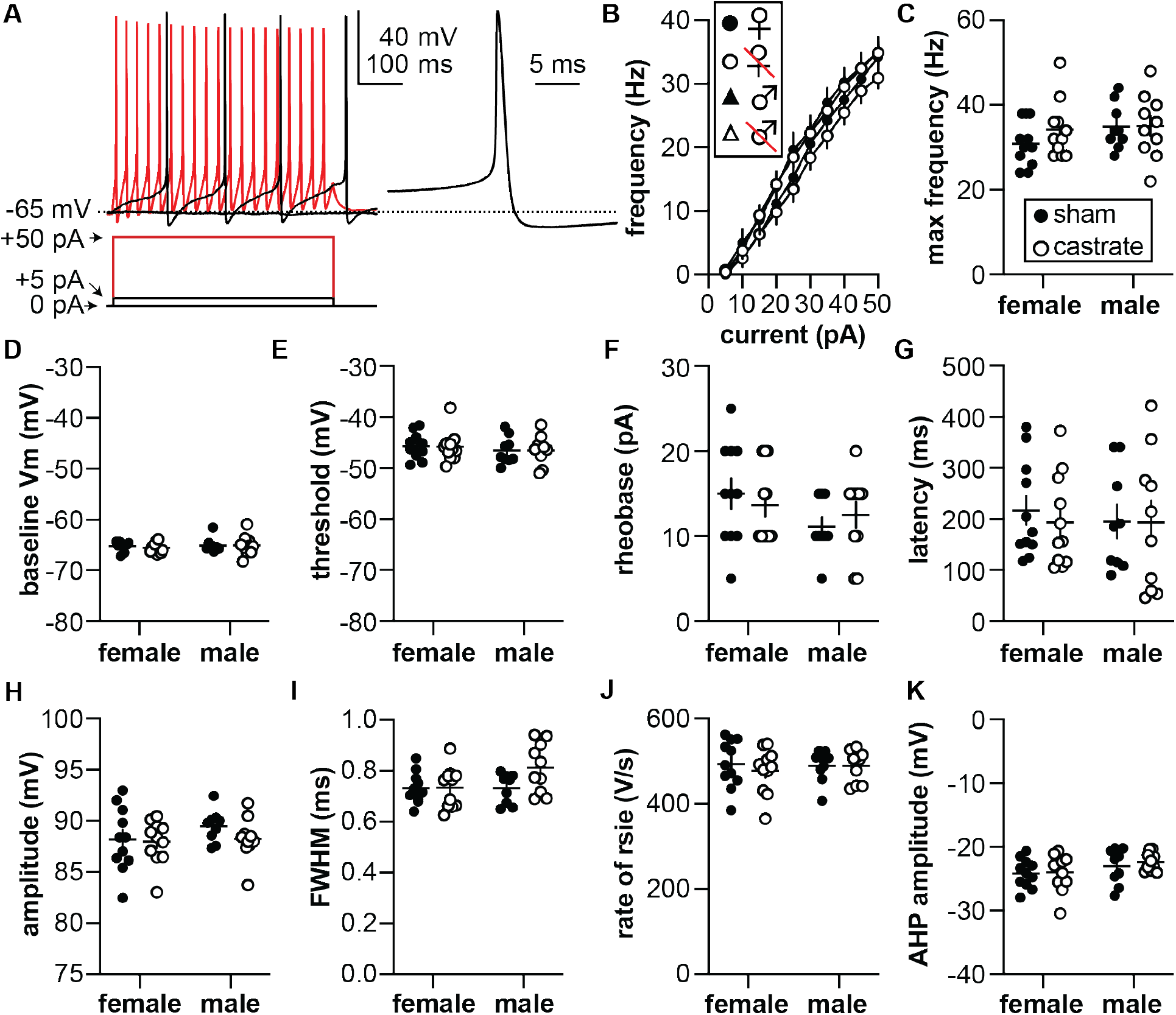
Action potential properties. (A) (left) Representative membrane voltage (top) responses to current injections (bottom); only three steps are shown for clarity. (Right) Expanded first action potential waveform. (B) Frequency of action potentials as a function of current injection. (C) Maximum action potential frequency. (D) Baseline membrane potential before rheobase current injection. (E) Action potential threshold. (F) Rheobase (minimum current to produce an action potential). (G) Action potential latency. (H) Action potential amplitude. (I) Action potential full width at half maximum (FWHM). (J) Maximum rate of rise of the action potential. (K) Amplitude of the afterhyperpolarization. Note y-axes of D, E, and H do not start or end at zero. Statistical parameters are in Tables 5 and 6.

## Discussion

A century ago, early studies of pituitary ablation and replacement revealed a link between the emerging science of neuroendocrinology and reproductive processes (Evans HM, 1921; Smith, 1927). Around the same time, the reverse link was being made by observing the effects of orchidectomy upon the pituitary, and amelioration of these effects by replacement with fat-soluble extracts of the testes (McCullagh, 1932). It is now well established that reproduction revolves around the stimulatory and feedback interactions of the hypothalamo-pituitary-gonadal axis, with GnRH neurons serving as the final common pathway for central signaling to the pituitary. These interactions are homeostatic in males and homeostatic throughout most of the reproductive cycle in females; removal of the gonads opens these homeostatic feedback loops and results in increased GnRH neuron activity, GnRH release and LH release in both sexes (Leipheimer et al., 1985; Levine et al., 1985; Karsch et al., 1987; Caraty and Locatelli, 1988; Condon et al., 1988; Jackson and Kuehl, 2000; Han et al., 2020). Despite the similarity in steady-state endocrine response, it is possible that latent sex differences in underlying neurobiological mechanisms exist (Jain et al., 2019). We tested the hypotheses that removal of negative homeostatic feedback would reduce potassium currents and increase the excitability of GnRH neurons. Based on the parameters we quantified, no substantial differences were observed that could be attributed to sex or feedback condition. We thus reject these hypotheses.

The present data showing no changes in the intrinsic properties of GnRH neurons in females between diestrous and OVX mice support prior work assessing excitability of GnRH neurons in OVX vs OVX+E mice exhibiting diurnal changes between negative and positive feedback (Christian et al., 2005). In this model, GnRH neuron activity and LH release are suppressed in the morning (AM) in OVX+E mice relative to OVX mice, demonstrating homeostatic negative feedback, but increased in the afternoon (PM) in cells from OVX+E mice, demonstrating estradiol positive feedback. Interestingly there were no differences in GnRH neuron excitability between OVX mice at either time of day and OVX+E mice in the AM (negative feedback), whereas GnRH neurons from OVX+E mice recorded in the PM during positive feedback were more excitable (Adams et al., 2018b). The present work extends these data to include both a comparison between the sexes and a comparison using the open loop OVX/ORX condition vs sham-operated controls to ameliorate caveats associated with steroid hormone replacement. The lack of effect of removing homeostatic feedback suggests a majority of the changes that lead to increased GnRH output in gonadectomized animals are processed presynaptic to these cells. In and of itself, this concept is not new as GnRH neurons do not appear to express detectable levels of steroid hormone receptors other than the beta isoform of the estradiol receptor (Herbison et al., 1996; Hrabovszky et al., 2000; Hrabovszky et al., 2007). There has thus long been a relative consensus in the field that steroid feedback is integrated via upstream, steroid-responsive cells (Wintermantel et al., 2006). The lack of observed changes in GnRH neuron properties examined suggests homeostatic negative feedback signals are conveyed from these afferents in a manner that alters the output of GnRH neurons without substantial biophysical changes at the level of the cell soma. Of note, there is increased frequency of GABA_A_-receptor-mediated postsynaptic currents, which can excite GnRH neurons (DeFazio et al., 2002; Herbison and Moenter, 2011), observed in GnRH neurons from ORX vs intact males (Chen and Moenter, 2009), and in cells from OVX vs OVX+E females during negative feedback (Christian and Moenter, 2007). Postsynaptic currents have relatively short-lived effects on membrane potential that may not lead to changes in overall excitability. Interestingly, changing homeostatic negative feedback does engage mechanisms affecting excitability in arcuate kisspeptin neurons, also known as KNDY neurons for their coexpression of three neuropeptides: kisspeptin, neurokinin B and dynorphin (Oakley et al., 2009; Moore et al., 2018). KNDy neurons have been postulated to be key afferent inputs to GnRH neurons to drive episodic release from these cells (Qiu et al., 2016; Clarkson et al., 2017; Vanacker et al., 2017). In KNDy neurons from OVX+E vs OVX females examined during negative feedback in the morning, estradiol reduced fast transient I_A_ (DeFazio et al., 2019), and I_A_ modifies action potential patterns in these cells (Mendonca et al., 2018).

The lack of changes in the homeostatic vs open loop models studied in the present work stand in marked contrast to changes that occur in GnRH neurons from females during estradiol positive feedback (Adams et al., 2018a; Adams et al., 2018b; Adams et al., 2019). In addition to the increase in GnRH neuron excitability observed during positive feedback mentioned above, GnRH neurons exhibit reduced transient potassium currents and increased high-voltage-activated calcium conductances in OVX+E mice during positive feedback (DeFazio and Moenter, 2002; Sun et al., 2010). While both the removal of negative feedback by gonadectomy and induction of positive feedback both increase GnRH release, the nature of this increase is quite different. Removal of negative feedback maintains the episodic GnRH/LH release pattern that is characteristic in males and most of the reproductive cycle in females (Leipheimer et al., 1985; Levine et al., 1985; Karsch et al., 1987; Caraty and Locatelli, 1988; Condon et al., 1988; Jackson and Kuehl, 2000; Han et al., 2020). Pulse frequency and amplitude are typically increased. Even in long-term castrated rams, when the pulsatile nature of LH release is no longer evident, high frequency GnRH pulses are clearly observed (Caraty and Locatelli, 1988). Induction of positive feedback, however, shifts the pattern from episodic to a continual elevation above baseline that lasts for several hours (Sarkar et al., 1976; Levine and Ramirez, 1982; Clarke et al., 1987; Moenter et al., 1990; Moenter et al., 1991; Moenter et al., 1992a; Moenter et al., 1992b; Xia et al., 1992). Given the fundamental differences in how the output of GnRH neurons is altered in these two circumstances, it is reasonable to postulate that more extensive changes are required for successful positive feedback, and that this is in part accomplished by extending the mechanisms engaged to the alteration of GnRH neuron intrinsic properties to allow continuous secretion to be maintained. Conceptually, pulse frequency can be rapidly modulated by homeostatic perturbances that influence reproduction. For example, inflammatory stress (Battaglia et al., 1998), hypoglycemia (Chen et al., 1996), and naloxone antagonism of opiates (Caraty et al., 1987) all rapidly induce changes in the pulse pattern of GnRH or multiunit activity associated with LH in the hypothalamus. In contrast, estradiol induction of positive feedback is a process with a substantial (typically >12h) delay to the increased release and this increase appears to be all or none as it is not dependent upon continued presence of estradiol (Evans et al., 1997).

The apparently different postsynaptic effects of removing negative feedback and inducing positive feedback raise some interesting questions for future contemplation. This is particularly true given that the prevailing view in the field is that the neuromodulator kisspeptin plays an important role in activating both episodic and surge modes of GnRH release (Porteous and Herbison, 2019; Wang et al., 2019). Kisspeptin application in brain slices reduces I_A_ in a manner similar to induction of estradiol positive feedback (Pielecka-Fortuna et al., 2011), and also increases excitability of GnRH neurons (Adams et al., 2018b), indicating it can alter the intrinsic properties of these cells. The two hypothalamic populations of kisspeptin neurons, the aforementioned KNDy neurons in the arcuate and those in the anteroventral periventricular area (AVPV) are thought to mediate pulsatile and surge modes of GnRH release, respectively. These populations possess different cotransmitters and mediators (Cravo et al., 2011; Skrapits et al., 2015); if co-released with kisspeptin these could produce counteracting effects in the postsynaptic GnRH neuron. Alternatively other afferent populations may have critical roles. In this regard, the increase in GnRH neuron firing induced by kisspeptin application is longer in duration than the typical pulse (Moenter et al., 1992b; Evans et al., 1996; Han et al., 2005; Pielecka-Fortuna et al., 2008). The neuromodulator known as gonadotropin-inhibitory hormone in birds and RFRP3 in mammals can counteract the activating effects of applied kisspeptin on GnRH neuron firing rate in brain slices by activating a potassium current (Wu et al., 2009). While speculative, an interplay of kisspeptin and RFRP3 may have opposing effects on intrinsic GnRH properties.

While we have confidence in the lack of effect of sex or removing homeostatic feedback on the properties examined in the present work, it is important to acknowledge alternative possibilities and mechanisms. It is possible that removing negative feedback induces equal and opposite changes in intrinsic properties resulting in masking of these changes in the current-clamp recordings. This seems unlikely given the different kinetic properties of the voltage-gated channels likely to mediate changes in action potential firing and properties. It is also possible that changes are induced in the biophysical properties of GnRH neurons that are distal to the soma and not possible to monitor in our brain slice preparation. For example, arcuate kisspeptin neurons can interact with GnRH neurons via kisspeptin and neurokinin B at the level of the terminals in the median eminence (Gaskins et al., 2013; Glanowska and Moenter, 2015; Yip et al., 2015). Examining these properties at a different point post gonadectomy may have revealed sex differences. While both males and females increase GnRH and LH release following gonadectomy, the increase is somewhat delayed in females. At the time point we investigated of 5-7 days post gonadectomy, clear increases in LH have occurred in both sexes (Yamamoto et al., 1970). It is also possible that removing homeostatic feedback alters other aspects of GnRH neurons to increase GnRH output, such as increasing GnRH mRNA (Finn et al., 1998; Gore, 1998), altering ionotropic receptors that produce brief changes in membrane potential, or changing excitation secretion coupling to make it more effective. Any of these alternative mechanisms could be sexually differentiated.

In sum, the present work rejected the hypotheses that removing homeostatic gonadal feedback induces changes in potassium currents and excitability of GnRH neurons. In so doing, we provide evidence for different mechanistic strategies to regulate the output of GnRH neurons during homeostatic vs positive feedback.

## Abbreviations

GnRH: gonadotropin-releasing hormone
LH: luteinizing hormone
4AP: 4-aminopyridine
OVX: ovariectomized
ORC: orchidectomized
GDX: gonadectomized
AHP: afterhyperpolarization
FWHM: full width at half maximum

## Acknowledgements

We thank Elizabeth Wagenmaker and Laura Burger for expert technical assistance and Joseph Starrett, Elizabeth Wagenmaker and Jennifer Jaime Alvarez for editorial comments. We thank James L. Kenyon, University of Nevada, Reno, for the Excel spreadsheet used to calculate junction potentials.

